# Polymorphic centromere locations in the pathogenic yeast *Candida parapsilosis*

**DOI:** 10.1101/2020.04.09.034512

**Authors:** Mihaela Ola, Caoimhe E. O’Brien, Aisling Y. Coughlan, Qinxi Ma, Paul D. Donovan, Kenneth H. Wolfe, Geraldine Butler

## Abstract

Centromeres pose an evolutionary paradox: strongly conserved in function, but rapidly changing in sequence and structure. However, in the absence of damage, centromere locations are usually conserved within a species. We report here that isolates of the pathogenic yeast species *Candida parapsilosis* exhibit within-species polymorphism for the location of centromeres on two of its eight chromosomes. Its old centromeres have an inverted-repeat (IR) structure, whereas its new centromeres have no obvious structural features, but are located within 30 kb of the old site. Centromeres can therefore move naturally from one chromosomal site to another, apparently spontaneously and in the absence of any significant changes in DNA sequence. Our observations are consistent with a model where all centromeres are genetically determined, such as by the presence of short or long IRs, or the ability to form cruciforms. We also find that centromeres have been hotspots for genomic rearrangements in the *C. parapsilosis* clade.

## INTRODUCTION

Centromeres are the point of assembly of the kinetochore, the position at which the spindle microtubules are connected to the chromosomes, enabling efficient and accurate separation of chromosome/chromatid pairs during cell division. Most eukaryotes have large “regional” centromeres that have been proposed to be epigenetically determined. They are specified by arrays of chromatin, compacted by di- or tri-methylation at lysine 9 of histone H3 (H3K9me2/3). The position of the centromere in most species is determined by the presence of a variant of histone H3, called CENP-A in mammals, or Cse4 in yeast.

Centromere repositioning occurs on an evolutionary time scale, leading to the formation of evolutionarily new centromeres (ENCs). ENCs have played an important role in speciation, including in many mammals (Rocchi et al. 2012; Stanyon et al. 2008). An ancient ENC at one chromosome in orangutans is polymorphic; individuals can be homozygous for either the old or the new centromere, or heterozygous for both (Locke et al. 2011; Rocchi et al. 2012). The new centromere location lacks the repetitive alpha satellites observed at other centromeres. In addition, damage to, or loss of, existing centromeres can be rescued by the formation of new (neo) centromeres at different locations. Neocentromere formation following damage has been observed in human clinical samples, as well as in other primates, in equidae, marsupials, plants and yeasts (reviewed in (Rocchi et al. 2012; Schubert 2018; Burrack and Berman 2012)). Movement of centromeres among individuals within a species in a non-clinical context is much more rarely described. A small number of neocentromeres formed in human cells that have no obvious clinical effect have been reported; these were usually observed during routine amniocentesis (reviewed in (Rocchi et al. 2012)). In addition, the location of one centromere in the horse (devoid of satellite DNA) varies among individuals (Purgato et al. 2015; Wade et al. 2009). The mechanisms underlying the formation of new centromeres are not fully understood, though many are likely to be associated with chromosomal inversion and translocation (Schubert 2018). The formation of neocentromeres following damage is particularly well studied in the yeast *Candida albicans* (Burrack and Berman 2012). Koren et al (Koren et al. 2010) suggested that in this species, centromeres are associated with the presence of early origins of replication, and that the formation of neocentromeres changes the activity of nearby origins.

Basic centromere organization is conserved in many fungi, including the basidiomycetes and the filamentous ascomycetes (Friedman and Freitag 2017). Centromeres in the budding yeasts (the Saccharomycotina) have undergone substantial changes associated with the loss of the lysine methylation machinery (Malik and Henikoff 2009). Within Saccharomycotina, the Saccharomycetaceae clade, containing the model yeast *Saccharomyces cerevisiae*, is by far the best studied. These species have small “point” centromeres, where function is determined by sequence. The *S. cerevisiae* centromere consists of three conserved regions called Centromere Determining Elements - CDEI, CDEII and CDEIII (Schulman and Bloom 1991). Cse4 is present in one nucleosome at the centromere (Meluh et al. 1998; Furuyama and Biggins 2007; Henikoff and Henikoff 2012). Similar point centromeres are found in other Saccharomycetaceae species (Kitada et al. 1997; Gordon et al. 2011; Mattei et al. 2002). In *Naumovozyma* species, the sequences of the CDE elements are different, but they still act as point centromeres (Kobayashi et al. 2015). The point centromeres in *S. cerevisiae* are among the fastest evolving sequences in the genome (Bensasson et al. 2008). However, point centromeres are not present in most fungal genomes (Malik and Henikoff 2009).

Centromere structure has also been investigated in other families in the Saccharomycotina, including the Pichiaceae and the CUG-Ser1 clade. Within the Pichiaceae, centromere structure is known in *Kuraishia capsulata* and *Komagataella phaffii*. In *K. capsulata*, centromeres lie in 2-6 kb regions with low GC content, and a 200 bp motif is conserved across some chromosomes (Morales et al. 2013). In *K. phaffii* the centromeres consist of a 1 kb central (mid) region, flanked by a 2 kb inverted repeat (IR) (Coughlan et al. 2016). There is no conservation in sequence among the four centromeres in *K. phaffii*, and Cse4 localizes across the mid region and the IR.

The CUG-Ser1 clade within the Saccharomycotina contains many *Candida* and other species, characterized by translating CUG as serine rather than leucine (Ohama et al. 1993). The centromeres of *Candida albicans* and *Candida dubliniensis* are described as “small regional”; they are characterized by gene-free regions of 4-18 kb, with 3-5 kb occupied by Cse4 (Sanyal et al. 2004; Roy and Sanyal 2011; Padmanabhan et al. 2008). The flanking compact chromatin extends up to 25 kb for *C. albicans* CEN7 (centromere of chromosome 7) (Sreekumar et al. 2019). There is no sequence conservation between centromeres of different chromosomes. There are short unique inverted repeats surrounding *C. albicans* CEN1, CEN4, and CENR, and longer repeats surrounding CEN5 (Sanyal et al. 2004). In the related species *Candida tropicalis*, the centromere cores are all flanked by IRs, and there is significant sequence conservation between different centromeres (Chatterjee et al. 2016). Centromeres in the more distantly related *Clavispora lusitaniae* have 4 kb regions occupied by Cse4, with no sequence conservation (Kapoor et al. 2015). The *C. lusitaniae* centromeres lie in regions with low GC content, which has also been proposed to mark centromeres in the CUG-Ser1 clade species *Debaryomyces hansenii* and *Scheffersomyces stipitis* (Lynch et al. 2010). The putative centromeres in these latter species contain clusters of retrotransposons (Lynch et al. 2010; Coughlan et al. 2016).

In this work, we experimentally determined the location of centromeres in the CUG-Ser1 clade yeast, *Candida parapsilosis*, and we inferred the locations in the related species *C. orthopsilosis* and *C. metapsilosis*. We identified *C. parapsilosis* centromeres by ChIP-Seq and we show that they usually have an IR structure. However, we also identified one *C. parapsilosis* isolate in which two centromeres have moved, indicating that centromere location in this species is polymorphic. The new centromere locations are <30 kb from the old ones, but they do not have any IR structure. These are the first examples of the birth of centromeres *de novo* in natural fungal isolates, and among the few observed in any species (Purgato et al. 2015; Wade et al. 2009; Locke et al. 2011; Rocchi et al. 2012). The new centromeres have no obvious structure, and we conclude that centromeres can change easily from IR-type to apparently epigenetic-type over short time scales, as well as over evolutionary time. In addition, we find that chromosomal rearrangements between *C. parapsilosis* and its closest relatives are enriched at centromeres, and that multiple rearrangements have occurred at, or very close to the centromeres, indicating fragility and dynamic evolution.

## RESULTS

### Identification of centromeres in *C. parapsilosis*

Many fungal centromeres are located in large intergenic regions and may be flanked by IR sequences. When we looked for regions that matched these criteria in the genome of *C. parapsilosis* CDC317 (the sequenced reference genome (Butler et al. 2009)) we identified one candidate centromere per chromosome (Fig. 1A,B). These regions range from 5.8 to 7.1 kb and lack genes. Each contains an inverted repeat (IR) sequence (shown in red in the dot matrix plot Fig. 1A), flanking a middle (mid) sequence. The IRs vary in size. Some are relatively short (e.g. 443 bp on chromosome 6) and in others the repeat region is broken into several sections (e.g. chromosome 1, total size ~ 1600 bp). The similarity between IRs ranges from 85 to 96.7%. Importantly, the sequence of the IRs are conserved among chromosomes, and the conservation extends beyond the IRs (Fig. 1B, black boxes). All IRs are predicted to form large secondary structures using RNAFold (Lorenz et al. 2011). However, there is no conservation among the mid regions that lie between the IRs on different chromosomes.

**Fig. 1.**
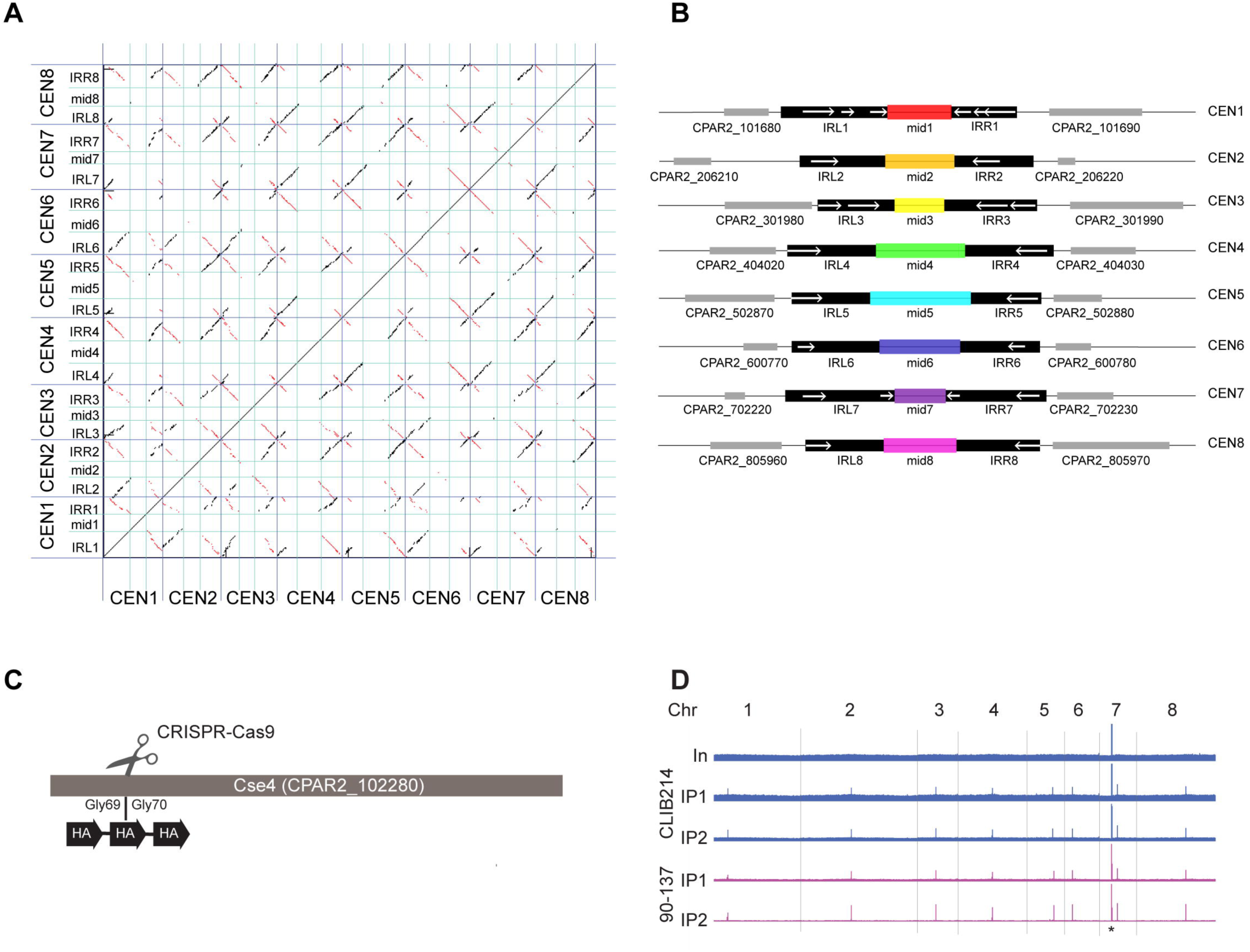
*C. parapsilosis* centromeres consist of unique mid regions surrounded by partially conserved Inverted Repeats. A. Dot matrix plot comparing the putative centromere sequences in *C. parapsilosis*. Centromere regions (Supplemental_Table_S2) were concatenated and are delineated by dark blue lines. Inverted repeats (Right, IRR and left, IRL) are separated with cyan lines. Each dot represents a 25-bp window. Inverted sequences are shown in red, and direct repeats in black. B. Diagrammatic representation of the information in (A). Regions that are conserved among chromosomes are shown in black. Locations of inverted repeats (> 75% DNA sequence identity) are shown with white arrows. The mid regions are illustrated in different colors that indicate that each of them has a unique sequence. Adjacent genes are shown in gray. Each region shown is approximately 10 kb in length. C. Three copies of an HA tag were introduced into both alleles of the endogenous *CSE4* gene in *C. parapsilosis* CLIB214 and 90-137 using CRISPR-Cas9 editing. The gene was cut between Glycine 69 and Glycine 70, and a repair template containing the HA tags was inserted by homologous recombination. The construct was confirmed by sequencing. D. Visualization of the ChIP-seq signal across all chromosomes (Chr) in Cse4-tagged derivatives of *C. parapsilosis* CLIB214 and 90-137. In = Input (before immunoprecipitation), IP1 and IP2 show two independent immunoprecipitation replicates from each strain. Strains derived from *C. parapsilosis* CLIB214 are shown in blue, and from 90-137 in purple. There is one signal per chromosome in the IP samples, identifying the centromere, except for chromosome 7, for which the rDNA locus (black asterisk) also generates a signal. The X-axis in each plot is the chromosome coordinates and the Y-axis is the number of reads mapping to a position. The maximum scale for *C. parapsilosis* CLIB214 is restricted to reduce the signal from the rDNA. Data is visualized using Integrated Genomics Viewer (IGV) (Thorvaldsdóttir et al. 2013).

To validate these predictions, we determined the location of the variant histone H3, Cse4, by chromatin immunoprecipitation (ChIP). *C. parapsilosis* has a diploid genome. We introduced three copies of a 9 amino acid epitope from human influenza hemagglutinin (HA), near the N terminus of both Cse4 alleles using CRISPR-Cas9 editing together with a synthetic repair template (Lombardi et al. 2017) (Fig. 1C). The epitope was introduced into Cse4 twice independently in two different strains – *C. parapsilosis* CLIB214, which is the type strain, and *C. parapsilosis* 90-137, originally isolated from orbital tissue (Tavanti et al. 2005) and which can be efficiently edited using CRISPR-Cas9 (Lombardi et al. 2017, Lombardi et al. 2019a). We confirmed that the tagged protein is expressed and that it does not interfere with growth of the tagged strains, and we used ChIP-PCR to show that Cse4 binding is enriched at the predicted CEN1 sequence (Fig. S1).

To identify all the regions in the genome where Cse4 binds, we combined chromatin immunoprecipitation with DNA sequencing (ChIP-seq). We obtained one very strong ChIP-seq signal per chromosome that was present in only the immunoprecipitated Cse4-HA strains, and not in the input chromatin (Fig. 1D). We also identified a signal from the ribosomal DNA on chromosome 7, an artifact due to the high copy number which is also present in the control sample. More detailed analysis shows that the Cse4 signals from *C. parapsilosis* CLIB214 correspond with the regions that were bioinformatically identified as centromeres (Fig. 2). The centromeres are in regions that are devoid of open reading frames, and are generally low in transcription (Fig. 2). Unlike *C. tropicalis* (Chatterjee et al. 2016) but similar to *K. phaffii* (Coughlan et al. 2016) Cse4 binding extends beyond the mid regions into the IRs, reducing in frequency toward the ends of the repeats.

**Fig. 2.**
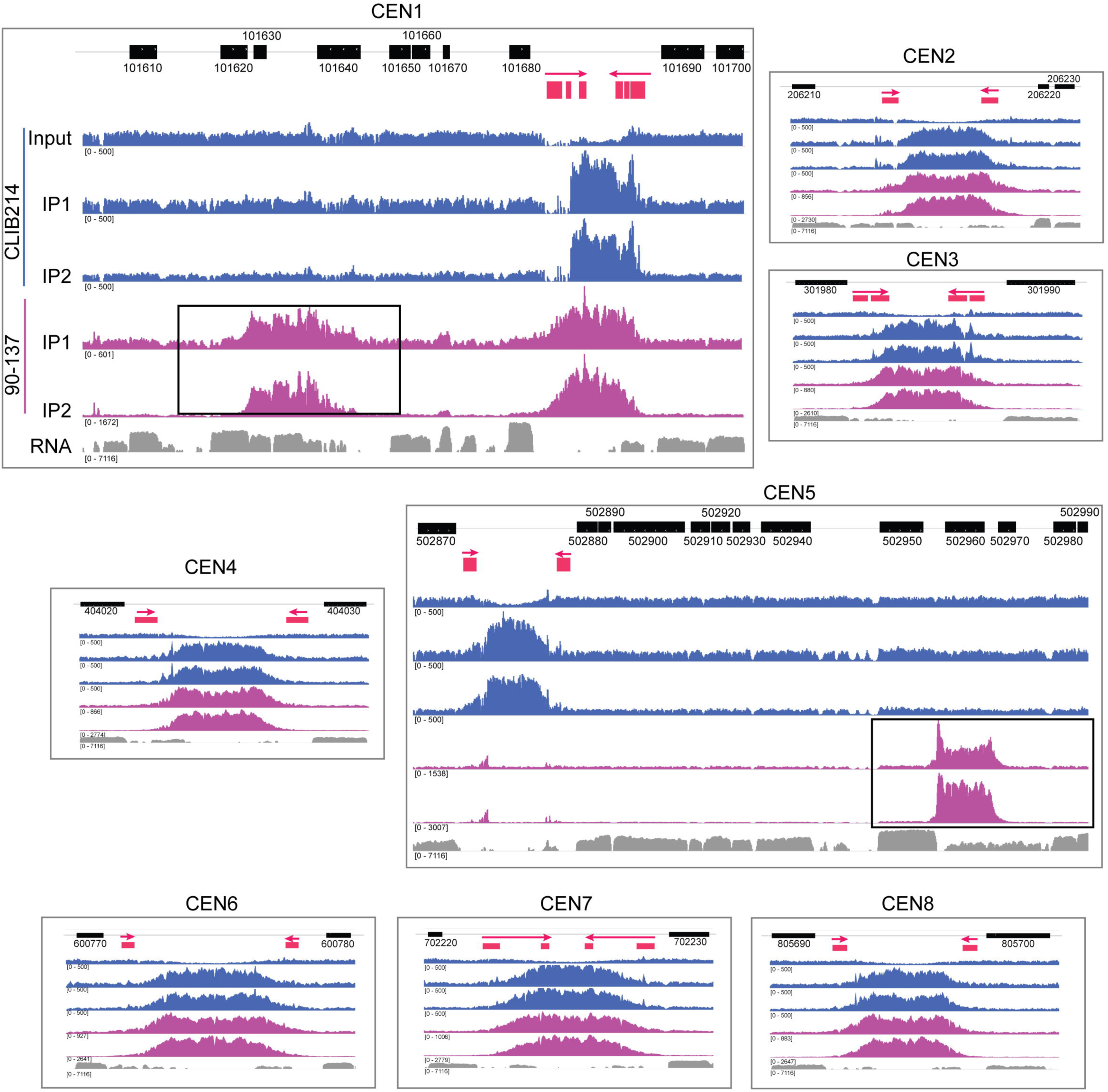
Natural polymorphisms for centromere location in *C. parapsilosis*. The ChIP-Seq data from Fig. 1D is shown in more detail, and the neocentromeres are highlighted with black boxes. The order of the tracks is the same in each panel, but is labelled for CEN1 only. The top track shows the location of *C. parapsilosis* protein coding genes. The second track shows the IR sequences only (red), with an arrow indicating the direction of the repeat. The extent of the regions conserved between chromosomes is not shown. ChIP-Seq read coverage is plotted in blue for *C. parapsilosis* CLIB214, and purple for *C. parapsilosis* 90-137. Two independent immunoprecipitation experiments were carried out per strain (IP1 and IP2). Only one control is shown; the total chromatin from *C. parapsilosis* CLIB214 (Input). The equivalent data for *C. parapsilosis* 90-137, and for an experiment with no tagged Cse4, is available at GEO, accession number GSE136854. The bottom track (gray) shows gene expression measured by RNA-seq during growth in YPD (taken from SRR6458364 from (Turner et al. 2018)). The read depth scale is indicated in brackets; the total number of reads varied in each experiment. The maximum scale for *C. parapsilosis* CLIB214 is restricted to 500 to reduce the signal from the rDNA. The RNA expression data is plotted on a log scale. The apparent dips in coverage at the centromeres in the input data is likely to be an artefact of the mapping procedure because reads that map to more than one site in the genome were discarded. Some reads are also incorrectly mapped to non-identical repeat sequences, resulting in a small Cse4 signal at CEN5 in 90-137. All data is visualized using IGV.

### Polymorphic centromere locations in *C. parapsilosis*

The Cse4 signal in *C. parapsilosis* 90-137 is very similar to *C. parapsilosis* CLIB214 (Fig. 1D). Closer examination shows the pattern is almost identical for 6 of the 8 chromosomes (Fig. 2). However, there are surprising differences at CEN1 and CEN5. For chromosome 1, there is a signal at the expected centromere in *C. parapsilosis* 90-137, similar to *C. parapsilosis* CLIB214. However, there is an additional signal, approximately 17 kb away in 90-137 (Fig. 2). This second signal, or neocentromere, partially overlaps two open reading frames, *CPAR2_101630* and *CPAR2_101640*, which are transcribed in *C. parapsilosis* CLIB214 (RNA track in Fig. 2). The difference is even more striking on chromosome 5. Here, *C. parapsilosis* 90-137 has no obvious Cse4 signal at the expected position of CEN5 (the small number of reads shown is an artefact of the mapping process, resulting from the presence of repeat sequences). Instead, the Cse4 signal is localized approximately 29 kb away, again overlapping transcribed ORFs, *CPAR2_502960* and *CPAR2_502970*. There are no IRs surrounding the new centromeres, and there is no sequence relationship with other centromeric regions.

We considered that the occurrence of neocentromeres in *C. parapsilosis* 90-137 might coincide with possible rearrangements of the chromosomes in this isolate. We therefore determined the genome structure of the Cse4-HA tagged strain using long read sequencing (Oxford Nanopore). The nuclear genome was assembled into 12 scaffolds > 1 kb in size (Fig. 3). The assembly failed at CEN6 on scaffolds 9/10. However, Fig. 3 shows that chromosomes 1 and 5 are collinear between *C. parapsilosis* 90-137/Cse4-HA and the reference genome, including around the centromere regions. The IR structures and mid region at the original CEN1 and CEN5 locations are intact in *C. parapsilosis* 90-137, and are 98 - 99% identical to the reference genome.

**Fig. 3.**
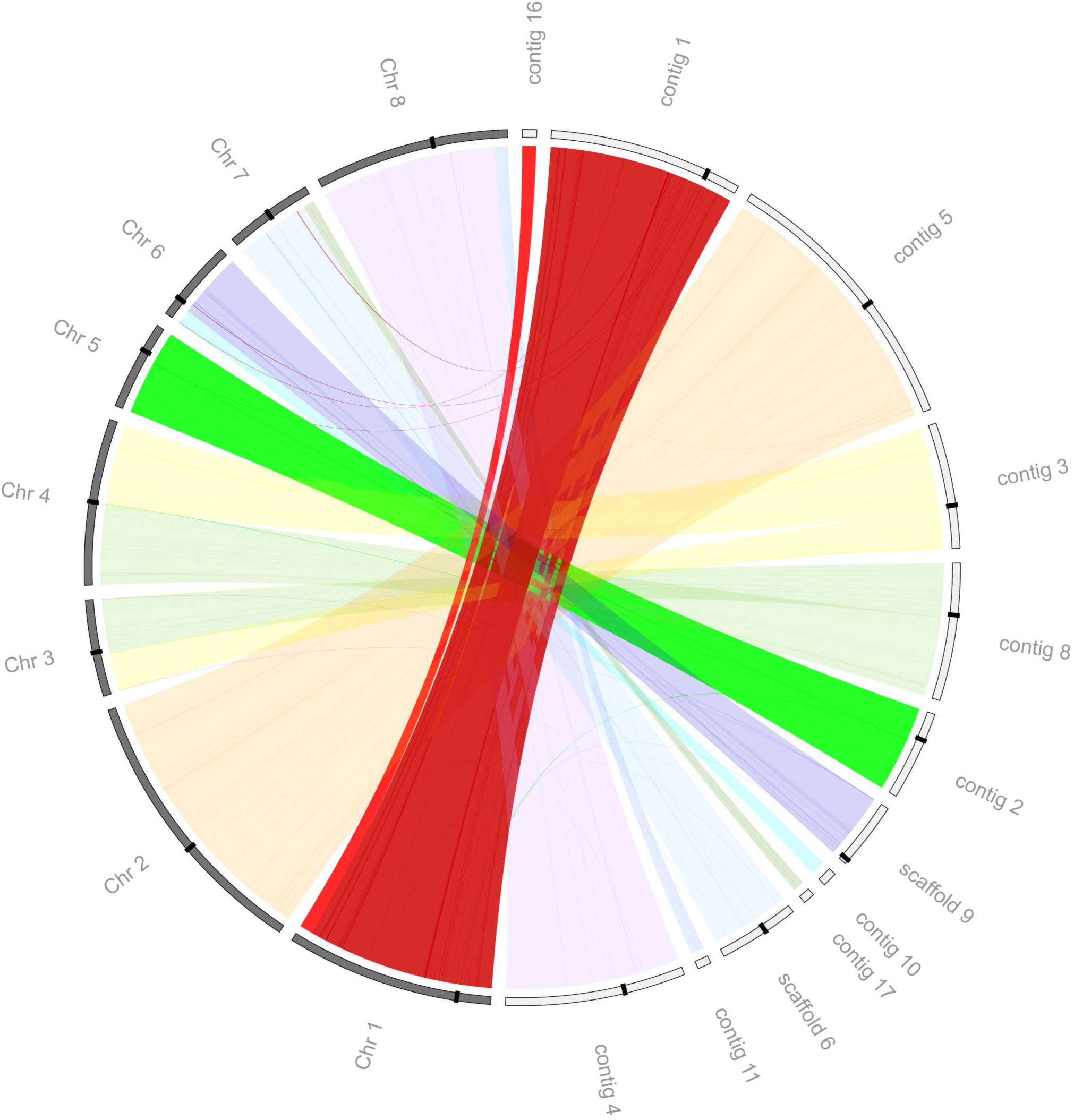
Lack of rearrangements at CEN1 and CEN5 in *C. parapsilosis* 90-137/Cse4-HA. The Circos plot compares the 8 chromosomes of the reference strain *C. parapsilosis* CDC317 (gray, left) to the 12 largest minION scaffolds from *C. parapsilosis* 90-137 (white, right). Centromeres are marked by black bands. Most chromosomes are collinear, including chromosome 1 (assembled in two contigs in 90-137, contig 1 and contig 16) and chromosome 5 (contig 2). There is an apparent translocation between chromosomes 3 and 4 (contig 3 and contig 8) at a repetitive gene that is near (but not at) the centromere. This may represent an error in the reference assembly, or a natural structural polymorphism.

The species *C. parapsilosis* is therefore polymorphic for centromere location on two chromosomes. The centromere relocations are associated with a transition from a structured (IR) format to a format with no obvious structure or sequence dependence, within a single species. On chromosome 5, it is likely that the centromeres on both copies of this chromosome have moved to a new location. It is possible that *C. parapsilosis* 90-137 is heterozygous at CEN1, with Cse4 at the expected location on one copy of chromosome 1 and at a new location on the other copy.

### Genomic rearrangements in *C. orthopsilosis* coincide with centromere locations

*C. parapsilosis* is closely related to *C. orthopsilosis* and *C. metapsilosis*; they are all members of the *C. parapsilosis sensu lato* clade (Tavanti et al. 2005). We surmised that the centromeres in these other species may have a similar structure to *C. parapsilosis*. The *C. orthopsilosis* 90-125 reference assembly (Riccombeni et al. 2012; Schröder et al. 2016) is not fully assembled at putative centromeres, so we used a minION assembly of this strain from Lombardi et al (Lombardi et al. 2019b). We identified one large region per chromosome likely to represent the centromere. The size of the regions ranges from 4.9 kb to 7.1 kb (Fig. 4A). Candidates on chromosomes 1, 2, 5, 6 and 7 have a similar structure to *C. parapsilosis* centromeres. A pair of IR sequences, varying in size from 788 bp on chromosome 5 to 2.2 kb on chromosome 6, flank a core region of ~ 3 kb. The similarity between IRs ranges from 91.0 to 99.8%, the sequences are conserved among chromosomes, and for chromosomes 5, 6, and 7 the conservation among chromosomes extends beyond the IRs. The remaining inferred centromeres (CEN3, 4, 8) do not contain IR sequences. However, 135 bp to 2.2 kb of the flanking regions surrounding the 2.6-3.4 kb mid regions are conserved with other centromeres. Like in *C. parapsilosis*, there is no conservation between the mid regions identified on different chromosomes. In addition, none of the *C. orthopsilosis* CEN regions (not just the IR-less ones) share significant sequence similarity with any of the *C. parapsilosis* CEN regions.

**Fig. 4.**
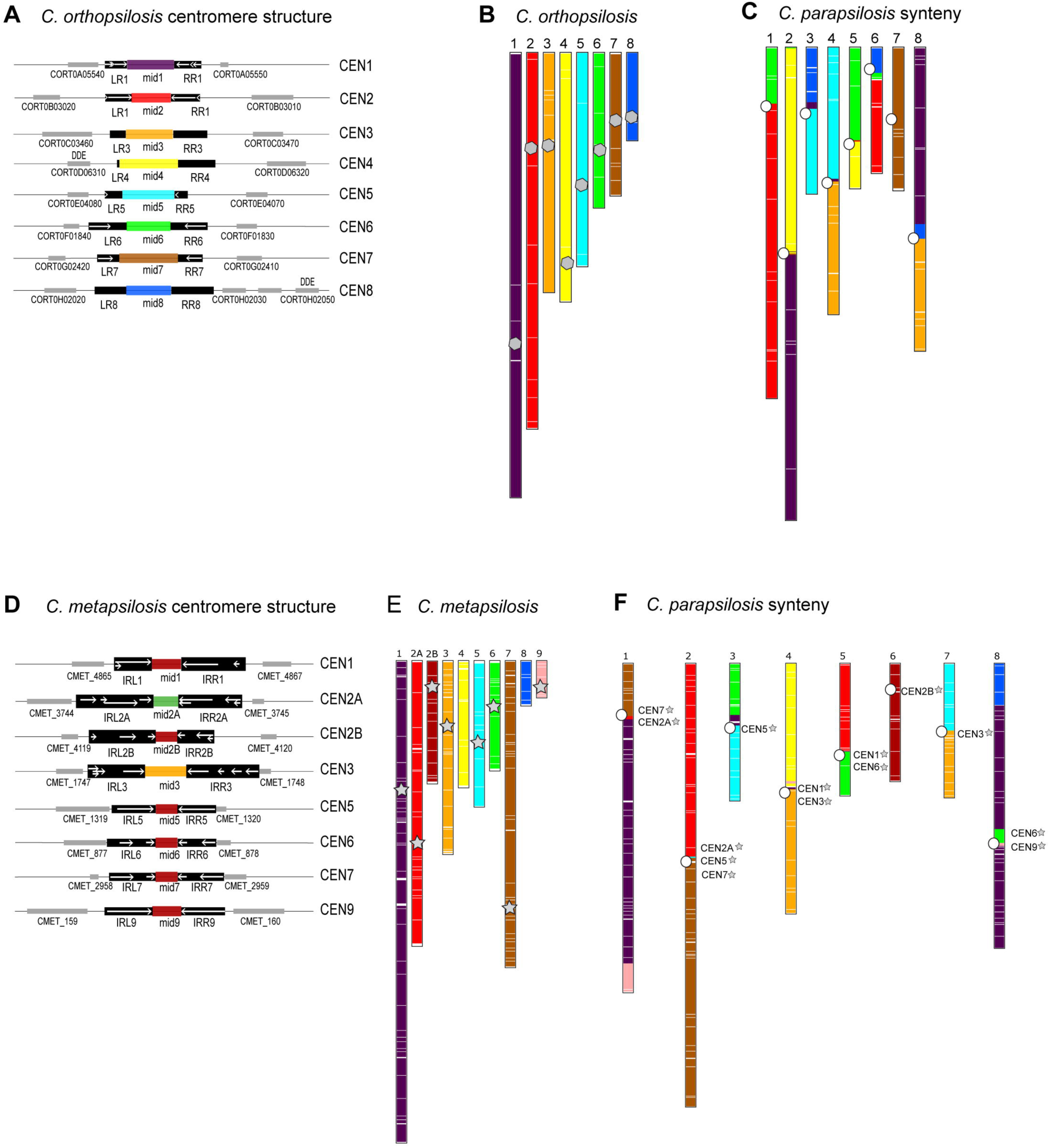
Identification of centromeres and centromere-proximal rearrangements in *C. orthopsilosis* and *C. metapsilosis*. A. Cartoon of centromere structure in *C. orthopsilosis* 90-125 (Lombardi et al. 2019b). All mid regions are unique and are shown in different colors. Sequences in black are conserved among chromosomes. IRs are shown with white arrows, and adjacent genes are shown with gray boxes. Putative transposases with DDE domains are indicated. More detail is provided in Supplemental_Fig_S2 and Supplemental_Table_S2. B and C. Synteny relationship between *C. parapsilosis* and *C. orthopsilosis*. SynChro (Drillon et al. 2014) was used (delta value of 2) to identify potential orthologs (reciprocal best hits, RBHs), represented by colored lines in the two species, and to generate synteny maps. B. Location of RBHs on *C. orthopsilosis* chromosomes. The approximate location of the putative centromeres are indicated with a gray polygon. C. *C. parapsilosis* chromosomes, colored with respect to the RBH from *C. orthopsilosis*. The location of the *C. parapsilosis* centromeres are indicated with an offset white circle. The location of syntenic *C. orthopsilosis* centromeres is shown in more detail in Fig. 5. D. Cartoon of centromere structure in *C. metapsilosis*. Sequences in black are conserved among chromosomes. IRs are shown with white arrows, which are sometimes fragmented and overlapping. Mid-core regions from some CENs are similar in sequence (>60%) and are shown in the same color. Adjacent genes are shown with gray boxes. More detail is provided in Supplemental_Fig_S2 and Supplemental_Table_S2. E and F. Synteny relationship between *C. parapsilosis* and *C. metapsilosis*. E. Location of RBHs on *C. metasilosis* chromosomes. The approximate location of the putative *C. metapsilosis* centromeres are indicated with a gray star (centromeres were not identified on scaffolds 4 and 8). F. *C. parapsilosis* chromosomes, colored with respect to the RBH from *C. metapsilosis*. The location of the *C. parapsilosis* centromeres are indicated with a white circle. The approximate location of syntenic *C. metapsilosis* centromeres are shown by name and with gray stars. The same colors are used for *C. orthopsilosis* (B) and *C. metapsilosis* (D). This does not indicate that synteny is completely conserved between these species; it is a feature of SynChro, which carries out pairwise comparisons.

We compared the conservation of centromere position and gene order between *C. parapsilosis* and *C. orthopsilosis* using SynChro, a tool designed to visualize synteny blocks in eukaryotic genomes (Drillon et al. 2014). Putative orthologs between the two species were assigned by identifying Reciprocal Best Hits (RBHs). Fig. 4B shows the locations of genes in *C. orthopsilosis* that have a RBH in *C. parapsilosis*. Each chromosome is assigned a specific color. Fig. 4C shows the locations of the same RBHs on the *C. parapsilosis* chromosomes, colored with respect to *C. orthopsilosis* chromosomes. It is immediately obvious that there is strong conservation of synteny between *C. orthopsilosis* and *C. parapsilosis*, as we have described previously (Riccombeni et al. 2012). One chromosome pair (chromosome 7 in each species) is essentially collinear, as shown by the brown color (Fig. 4B,C). Most of the other chromosomes are represented by two major colors in *C. parapsilosis*, indicating that there has been one major translocation per chromosome between *C. parapsilosis* and *C. orthopsilosis*.

Overlaying the position of the mapped centromeres shows that most of the evolutionary rearrangements between *C. parapsilosis* and *C. orthopsilosis* involve breakpoints at or near the *C. parapsilosis* centromeres (Fig. 4C). For some chromosomes there is a single breakpoint (e.g. chromosome 1). For others, whereas most of the two arms of the *C. parapsilosis* chromosome matches two *C. orthopsilosis* chromosomes, the junction near the centromere includes short sections from a third chromosome (e.g. on chromosome 8). These relationships are explored in Fig. 5, which shows the gene order around each *C. parapsilosis* centromere in more detail. Individual RBHs (identified and visualized using SynChro (Drillon et al. 2014)) are shown. Each *C. parapsilosis* centromere is compared to all *C. orthopsilosis* chromosomes, and syntenic blocks are highlighted.

**Fig. 5.**
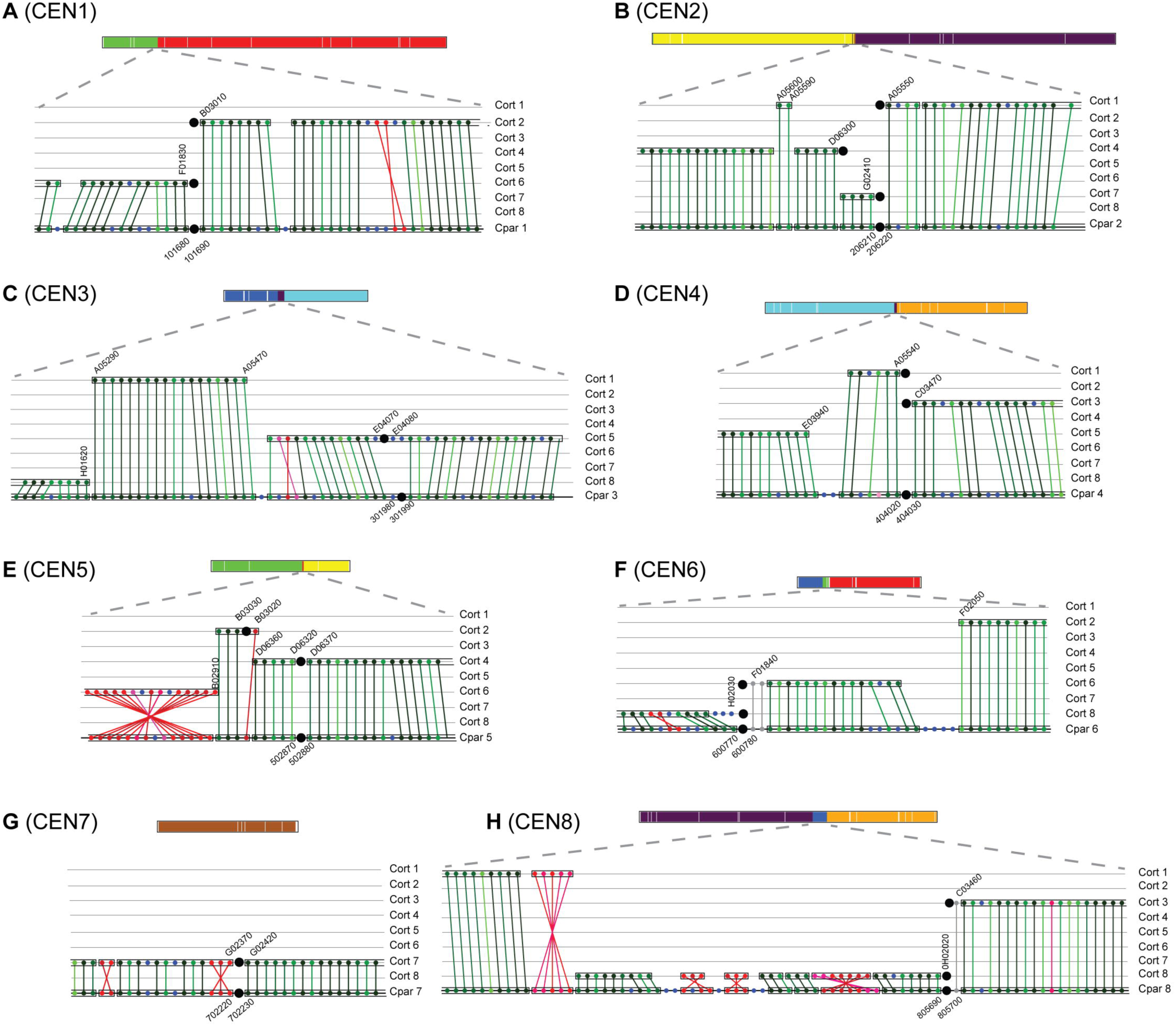
Interspecies synteny breakpoints occur at centromeres. Synteny between *C. parapsilosis* and *C. orthopsilosis* was visualized using SynChro (Drillon et al. 2014), with a delta value of 2. Changing delta values had minor effects on predicted synteny. A diagrammatic representation of each *C. parapsilosis* chromosome, colored as in Fig. 4C, is shown to scale at the top of each panel. The lower sections of each panel show the gene order around the centromere. A-F. Gene order around the 8 centromeres in *C. parapsilosis* compared to *C. orthopsilosis*. The bottom row in each panel shows gene order on the *C. parapsilosis* chromosome, and the 8 *C. orthopsilosis* chromosomes are shown above. Each gene is indicated by a colored dot, and RBH are joined by lines. Syntenic blocks are surrounded with a box. Centromeres are shown by large black circles. The chromosome number is indicated at the side of each panel. The names of some genes are shown for orientation purposes. The prefix “CORT0” has been removed from *C. orthopsilosis* genes and “CPAR2_” from *C. parapsilosis* genes for brevity. The color of the dots indicates the similarity of the proteins. Non inverted RBH are shown in green, ranging from darkest (>90% similarity) to lightest (<30% similarity), and inverted orthologs are shown in red. Genes without RBH orthologs are shown in blue. Genes in gray were not identified as RBH by SynChro, but were identified using CGOB (Maguire et al. 2013; Fitzpatrick et al. 2010).

Multiple rearrangements have occurred exactly at, or very close to, the centromere on almost all chromosomes (Fig. 5). For example, on *C. parapsilosis* chromosome 1, genes to the right of the centromere are syntenic with genes on *C. orthopsilosis* chromosome 2, and genes to the left of the centromere are syntenic with *C. orthopsilosis* chromosome 6 (Fig. 5A). The break in synteny coincides exactly with the location of the predicted centromeres on the two *C. orthopsilosis* chromosomes and with *C. parapsilosis* CEN1. More complex rearrangements are seen at CEN2, CEN4, CEN6, and CEN8 (Fig. 5). In each of these examples, there is a break in synteny at the *C. parapsilosis* centromere, so that the left and right flanks of the *C. parapsilosis* centromeres match two different *C. orthopsilosis* chromosomes, and the breakpoints in *C. orthopsilosis* also occur at or near its centromeres. However, in these four cases there are also additional rearrangements nearby, which at CEN2 (Fig. 5B) corresponds with a third centromere in *C. orthopsilosis* on chromosome 4.

Even on chromosome 7 (Fig. 5G), which is almost collinear between the two species, there has been an inversion beside the centromere. *C. parapsilosis* CEN3 is also collinear with *C. orthopsilosis* CEN5 (Fig. 5C). However, there have been two rearrangements on the left of *C. parapsilosis* CEN3, where a short block of genes on *C. parapsilosis* chromosome 3 matches a region on *C. orthopsilosis* chromosome 1. Most of the remainder of the left side of *C. parapsilosis* chromosome 3 is syntenic with *C. orthopsilosis* chromosome 6. Something similar is seen at *C. parapsilosis* chromosome 5 (Fig. 5E), except here one rearrangement occurs at a second *C. orthopsilosis* centromere (CEN2). In summary, *C. parapsilosis* has synteny breakpoints relative to *C. orthopsilosis* at 7 of its 8 centromeres, and most of these breakpoints also map to *C. orthopsilosis* centromeres. We examined the sequences around each interchromosomal rearrangement site but did not find any sequence repeats that could have facilitated the rearrangements.

### Genomic rearrangements in *C. metapsilosis* and *L. elongisporus*

*C. metapsilosis* originated from hybridization between two related species, generating a hybrid with a highly heterozygous diploid genome (Pryszcz et al. 2015). The best assembly of its genome is derived from Illumina sequencing only and is a consensus built from both haplotypes from two different isolates (Pryszcz et al. 2015). Of the nine largest *C. metapsilosis* scaffolds, we identified putative centromeres on seven (Fig. 4D,E). Scaffold 2 contained two candidate regions. Closer examination revealed that this scaffold contains a region (around *CMET_4044*) that is syntenic with two telomeres in *C. parapsilosis* (chromosomes 5 and 6). We do not know if this represents a recent telomere-to-telomere fusion in *C. metapsilosis* or if it is an assembly error. We split scaffold 2 at *CMET_4044*, generating scaffolds 2A and 2B (Fig. 4E), giving a total of eight centromeres. All the centromeres are surrounded by IRs, which have high levels of sequence similarity among chromosomes. The IRs on scaffolds 5, 6, 7 and 9 are relatively long (2.1-2.6 kb). IRs in conserved regions on scaffolds 6 and 7 are fragmented (Fig. 4D). IRs on scaffolds 1, 2A, 2B, and 3 are highly repetitive, with regions that sometimes overlap. The mid regions of *C. metapsilosis* centromeres vary in size from 1.2 to 2.2 kb, and unlike *C. parapsilosis* and *C.orthopsilosis*, there is sequence conservation among chromosomes. CEN2B, 5, 6, 7 and 9 share >75% identity, and CEN1 is approximately 60% identical to these (Fig. 4D, Supplemental_Fig. S2).

Fig. 4F shows a pattern of interspecies chromosomal breakage at centromeres between *C. metapsilosis* and *C. parapsilosis*, similar to that seen with *C. orthopsilosis*, although the rearrangements are different and have therefore occurred independently. *C. parapsilosis* chromosome 6 and *C. metapsilosis* scaffold 2B are collinear. Most other chromosomes have undergone a major rearrangement at points that correspond to the centromeres of both species. There have been complex rearrangements at these sites, similar to the *C. orthopsilosis*/*C. parapsilosis* comparisons. For example, the region around *C. parapsilosis* CEN2 is syntenic with regions near *C. metapsilosis* CEN2A, CEN5 and CEN7. Other apparent rearrangements may reflect gaps in the *C. metapsilosis* assembly (for example *C. metapsilosis* scaffold 8, which does not contain a centromere, maps to the end of *C. parapsilosis* chromosome 8).

*Lodderomyces elongisporus* is an outgroup to the *C. parapsilosis sensu lato* species group (Fitzpatrick et al. 2006). We did not find any structures similar to the *C. parapsilosis* centromeres in the *L. elongisporus* genome (Butler et al. 2009). However, Koren et al (Koren et al. 2010) hypothesised that centromeres in *L. elongisporus* are adjacent to early-firing origins of replication, as in *C. albicans*. They identified putative regions by characterizing GC skew, which switches between strands at replication origins. Koren et al (Koren et al. 2010) identified nine candidate centromeres in the 11 largest *L. elongisporus* scaffolds, that lie within intergenic regions and have a strong GC skew. Three may not represent true centromeres; one (on scaffold 9) is adjacent to the rDNA locus (Donovan et al. 2016) and two are in strongly transcribed regions (scaffold 7, scaffold 10) (Donovan et al. 2016) that are probably incorrectly annotated in the *L. elongisporus* genome. The most likely centromeres and a comparison of the synteny of *C. parapsilosis* with *L. elongisporus* are shown in Supplemental_Figure_S3. There are more rearrangements than observed between *C. parapsilosis* and *C. orthopsilosis* or *C. metapsilosis*. However, *C. parapsilosis* chromosome 6 and *L. elongisporus* chromosome 7 are collinear, and major rearrangements in the other chromosomes coincide with the location of the centromeres in *C. parapsilosis* and several of the remaining centromeres in *L. elongisporus* (Supplemental_Fig._S3). It is therefore likely that six of the proposed centromere locations in *L. elongisporus* are correct, and that centromeres are fragile sites in all four species. However, centromere structure in *L. elongisporus* is very different to the *C. parapsilosis sensu lato* species. There are no IRs, and the sequences are mostly unique (Koren et al. 2010). They are therefore more similar to the epigenetic centromeres described in *C. albicans* and *C. dubliniensis* (Thakur and Sanyal 2013; Sanyal et al. 2004; Padmanabhan et al. 2008).

In order to identify the number of translocations that have occurred during the evolution of the *C. parapsilosis* clade, we inferred the most likely ancestral chromosomal structure using AnChro (Vakirlis et al. 2016) (Supplemental_Fig_S4). Some of the reference assemblies are quite fragmented, and the number of predicted chromosomes in the ancestral species are probably overestimated (13-15, Supplemental_Fig _S4). It is therefore difficult to fully resolve every rearrangement. However, the synteny comparisons identified 13 interchromosomal breaks between *C. parapsilosis* and *C. orthopsilosis*, and all are at or close to the centromeres as shown in Fig 5. Most rearrangements occurred on the branch leading to *C. orthopsilosis* (Supplemental_Fig_S4). It is therefore clear that interchromosomal breaks are enriched at centromeres.

## DISCUSSION

Centromeres evolve remarkably rapidly, considering their conserved function (Henikoff et al. 2001). Species in the CUG-Ser1 clade have a very wide range of centromere types (Fig. 6). Centromeres of *C. albicans* and *C. dubliniensis* have been proposed to be epigenetically determined, and have little obvious sequence similarity, and few IRs (Sanyal et al. 2004; Padmanabhan et al. 2008). We have shown that the centromeres in the *C. parapsilosis sensu lato* species group consist of a mid region that is mostly unique, and is usually surrounded by IR sequences. The centromere structures in the *C. parapsilosis sensu lato* clade are most similar to those of *C. tropicalis* (Fig. 6; (Chatterjee et al. 2016; Padmanabhan et al. 2008)). However, in *C. tropicalis*, the mid regions of all centromeres are similar (~80% identity), and the IRs are highly homogenised. Chatterjee et al (Chatterjee et al. 2016) suggested that the ancestral centromere in *Candida* species consisted of an IR surrounding a core, and that most of the IRs have been lost in *C. albicans* and *C. dubliniensis*. Orthology of the centromeres on each chromosome within the CUG-Ser1 clade, despite their structural variation, is supported by evidence that gene order is partially conserved around centromeres among *C. albicans*, *C. dubliniensis* and *C. tropicalis* (Padmanabhan et al. 2008; Chatterjee et al. 2016). Synteny is conserved between *C. albicans* CEN3 and *C. parapsilosis* CEN5, and there is partial conservation of synteny around *C. albicans* CEN5 with centromeres in *C. parapsilosis*, *S. stipitis* and *C. lusitaniae*, even though centromeres do not contain IRs in the latter two species (Lynch et al. 2010; Chatterjee et al. 2016).

**Fig. 6.**
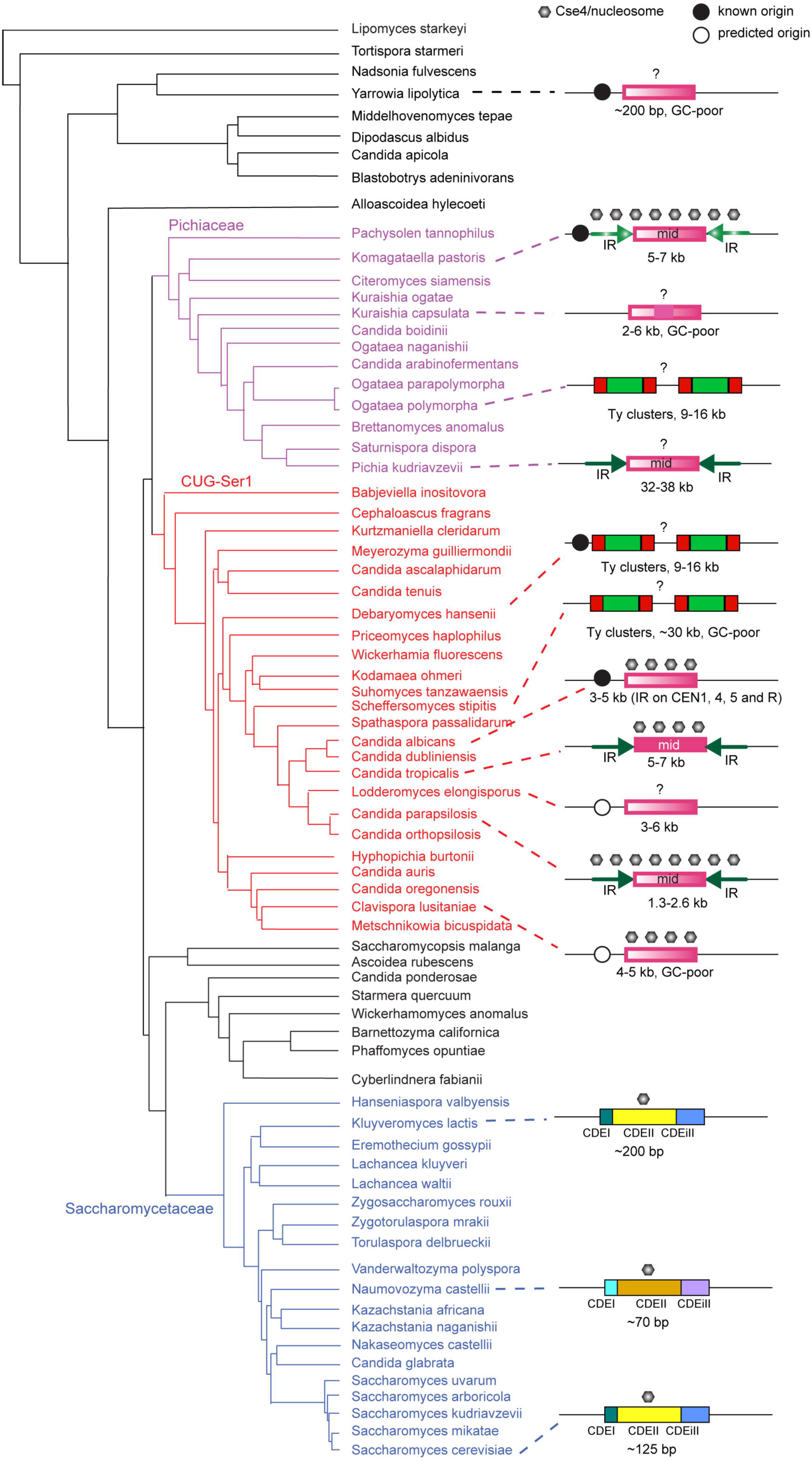
Organization of centromeres in Saccharomycotina species. The phylogeny is adapted from Shen et al (Shen et al. 2018). The size indicated on the centromeres refers to the region bound by Cse4 where known, or else where predicted bioinformatically, except for the Saccharomycetaceae where the size of the point centromere is shown. Solid color indicates conservation of sequence across centromeres in the same species, whereas a color gradient indicates unique sequences. Inverted repeats (IR) are shown with arrows, and Ty clusters as red and green boxes. Black circles show known (solid) or predicted (open) early firing origins of replication (see text for details). Point centromeres are conserved across the Saccharomycetaceae except for the Naumovozyma lineage, which have different sequences. Question marks indicate that localization of Cse4 nucleosomes has not been determined.

The IR structure of centromeres is likely to be old, because it is also found in some species in the sister clade, the family Pichiaceae (Fig. 6). In *Pichia kudriavzevii*, the IRs at each CEN are very similar and they are conserved across centromeres. In addition, these IRs share some similarity with mid sequences on other chromosomes (Douglass et al. 2018). In *Komagataella phaffii* (*Pichia pastoris*) both the IRs and the mid regions are unique at each CEN (Coughlan et al. 2016). The ancestor of the Pichiaceae and the CUG-Ser1 clade species therefore likely had an IR surrounding a mid region, with unique sequences at each centromere. The IRs have undergone homogenization in several species (*P. kudriavzevii*, *C. tropicalis* and *C. parapsilosis sensu lato*), and the mid regions have been homogenized in *C. tropicalis* and to a lesser extent in *C. metapsilosis*. IRs have probably been lost in *C. albicans*, *C. lusitaniae* and *K. capsulata*. In other species in the CUG-Ser1 clade (*D. hansenii*, *S. stipitis*) and in the Pichiaceae (*Ogataea polymorpha*) the CENs are associated with retrotransposons (Ty5-like elements). A retrotransposon (member of the Ty3/Gypsy family) is found at CEN7 in *C. tropicalis, C. albicans* and *C. dubliniensis* (Padmanabhan et al. 2008; Chatterjee et al. 2016). DDE-type transposases are found adjacent to *C. orthopsilosis* CEN4 and CEN8, but these are likely to be DNA transposons (Nesmelova and Hackett 2010), more similar to CEN-associated transposons in the basidiomycete *Cryptococcus neoformans* (Janbon et al. 2014).

It is not clear what the ancestral centromere structure was in the subphylum Saccharomycotina because centromeres have been characterized in very few species outside the Pichiaceae and the CUG-Ser1 clade (Fig. 6). The point centromeres in the Saccharomycetaceae are unusual, and probably represent a derived state (Lefrançois et al. 2013; Malik and Henikoff 2009; Kobayashi et al. 2015). Centromeric regions have been identified in *Yarrowia lipolytica*, an outgroup to the three clades (Fig. 6). These lie in regions of poor GC-content, adjacent to autonomously replicating sequences (Lynch et al. 2010; Fournier et al. 1993). *Y. lipolytica* centromeres may be small, and have conserved short palindromic repeats of 17 to 21 bp (Yamane et al. 2008). However, the exact structure of the centromere and the location of CENP-A (Cse4) in *Y. lipolytica* has never been determined. More experimental analysis of centromeres from other clades of the Saccharomycotina is therefore required before conclusions can be drawn about the ancestral centromere structure.

Kasinathan and Henikoff (Kasinathan and Henikoff 2018) postulated that all centromeres, whether apparently epigenetic or sequence-dependent, share a common feature – they are at regions that can make non-B form DNA. This can be achieved via dyad symmetry (IRs) in the DNA, or by the activity of specific DNA-binding proteins (such as binding of Cbf1 in the Saccharomycetaceae). IRs have the capacity to form cruciform structures, especially when associated with replication origins (Pearson et al. 1996). In particular, Kasinathan and Henikoff (Kasinathan and Henikoff 2018) found that neocentromeres in vertebrates are particularly enriched in regions of short dyad symmetry.

The formation of “rescue” neocentromeres when the endogenous centromere is damaged has been well studied in *C. albicans* (reviewed in (Burrack and Berman 2012)). When CEN5 or CEN7 is damaged, neocentromeres form, either adjacent to the original centromere or up to 450 kb away (Ketel et al. 2009; Thakur and Sanyal 2013). Koren et al (Koren et al. 2010) found that natural *C. albicans* CENs are near early firing replication origins, and that the formation of neocentromeres changes the timing of firing at adjacent origins. By characterizing the switches in base composition skew that occur at replication origins, they predicted that CENs are also near early firing origins of replication in *L. elongisporus*, *C. lusitaniae*, and *Y. lipolytica* (experimentally confirmed by Fournier et al (Fournier et al. 1993) for *Y. lipolytica*).

Examination of the known and predicted centromeres in CUG-Ser1 clade species shows that they all contain IRs (either long or short, including retrotransposon LTRs), and/or they are located near early firing replication origins (known or predicted). All of these structures can form cruciforms, which may be necessary to recruit Cse4, as has been reported for *Schizosaccharomyces pombe* (Folco et al. 2008). The loss of the IRs at centromeres in *L. elongisporus*, *C. lusitaniae*, and from some centromeres in *C. albicans* and *C. dubliniensis*, may be compensated by the presence of a nearby early-firing replication origin (Fig. 6). Therefore, there may be no true “epigenetic” centromeres in this clade; as Kasinathan and Henikoff (Kasinathan and Henikoff 2018) suggest, at least some part of centromere formation always requires cruciform or non-B form DNA, however it is made. The neocentromeres formed in *C. parapsilosis* 90-137 do not contain large IRs like the originals in this species. The hypothesis predicts that the neocentromeres form in regions capable of making cruciform structures, which may be facilitated by transcription. The *C. parapsilosis* neocentromeres are formed at regions that are transcribed, and transcription is known to facilitate centromere activity in *S. cerevisiae* (Ohkuni and Kitagawa 2011).

We found that the majority of chromosomal rearrangements between species in the *C. parapsilosis*/*L. elongisporus* clade involve breakpoints at or near centromeres, and that in several cases multiple closely-spaced breaks occurred near centromeres. Rearrangements between *C. albicans* and *C. tropicalis* also appear to be enriched around centromeres, which Chaterjee et al. (Chatterjee et al. 2016) suggested was facilitated by repeat sequences. However, rearrangements at centromeres in other species are unusual, and for example were rarely seen in Saccharomycetaceae species (Gordon et al. 2011; Dujon et al. 2004; Vakirlis et al. 2016). It therefore appears that centromeres are hotspots for chromosome breakage in the CUG-Ser1 clade, and particularly in species closely related to *C. parapsilosis* (for example, CENs in *C. albicans* and *C. dubliniensis* are collinear (Padmanabhan et al. 2008)). Although fragility may be associated with the presence of repeats (IRs) at the centromeres, and with the similarity of centromere sequences among chromosomes, even the centromeres of *L. elongisporus*, which have no IRs or other repeats, coincide with evolutionary breakpoints (Supplemental_Fig._S3). Interspecies rearrangements of the karyotype by breakage at centromeres has also been reported in the basidiomycete yeast *Cryptococcus* (Sun et al. 2017).

There are many unanswered questions about how and why the centromere relocations in *C. parapsilosis* 90-137 occurred. We do not know how frequent centromere location polymorphism is in *C. parapsilosis*, but the fact that we observed it in one of only two strains tested, affecting two of eight chromosomes, suggests that it is not rare. Such centromere sliding may also be frequent in other organisms (including humans), but has not been observed because of a lack of investigation (Rocchi et al. 2012). We also do not know what factors caused the original centromere sites to become disused in *C. parapsilosis* 90-137.

The IR structure at the original sites appears to be intact, so it is unclear why neither allele of CEN5 binds Cse4. Similarly, we do not know what makes the new centromere sites, at both CEN1 and CEN5, attractive for Cse4 binding. They have no repeats and no obvious features such as strong base composition skew. However, they are both within 30 kb of the original site, which means that diploids heterozygous for Cse4 bound at old and new sites (like at CEN1 in *C. parapsilosis* 90-137) can still establish proper spindle tension. Similar heterozygous centromeric sites have been reported in orangutans (Locke et al. 2011), horses (Purgato et al. 2015; Wade et al. 2009), and in *C. albicans* following damage at one allele (Thakur and Sanyal 2013). Lastly, we do not know why the new sites only bind Cse4 in *C. parapsilosis* 90-137 and not in *C. parapsilosis* CLIB214. Our discovery of “natural” neocentromeres in *C. parapsilosis* is one of the few known examples of within-species polymorphism for CEN locations, and provides an ideal opportunity for further future investigation of how centromere location and function is determined (Wade et al. 2009; Locke et al. 2011; Rocchi et al. 2012).

## METHODS

### Bioinformatic prediction of centromere location

Genomic sequences of intergenic regions larger than 2 kb were extracted from the reference sequence of *C. parapsilosis* CDC317 (Butler et al. 2009), *C. orthopsilosis* 90-125 (Riccombeni et al. 2012; Schröder et al. 2016), and the chimeric reference assembly of *C. metapsilosis* strains PL429 (SZMC1548) and SZMC8094 (Pryszcz et al. 2015) using a custom script (https://doi.org/10.6084/m9.figshare.9750638.v1). Sequences were compared using Blastn v 2.2.26 with default parameters and tabular alignment output (Altschul 1990). An IR pair was defined as a sequence identity >75% with a region in the opposite orientation (E-value cutoff 0.005). Candidate regions were selected for manual investigation. Predicted centromere locations in the *C. orthopsilosis* 90-125 reference assembly (Riccombeni et al. 2012; Schröder et al. 2016), available at CGOB (Fitzpatrick et al. 2010), had long regions of ambiguous bases, so we extracted equivalent regions from a minION assembly from Lombardi et al (Lombardi et al. 2019b) (Supplemental_Table_S2). Dot matrix plots were constructed using DNAMAN (www.lynnon.com) with a criterion of 23 matches per 25 bp window. Synteny was visualized using SynChro with a delta value of 2 (Drillon et al. 2014), using genome assemblies and annotations from CGOB (Fitzpatrick et al. 2010; Maguire et al. 2013). To reconstruct ancestral genomes, SynChro was run using delta values between 1 and 6. The ancestor of *C. parapsilosis* and *C. orthopsilosis* (A1) was reconstructed using AnChro (Vakirlis et al. 2016), varying delta values from 1 to 6 for each branch. *C. metapsilosis* and *L. elongisporus* were used as outgroups. The best A1 candidate, with the smallest number of chromosomes (13) and conflicts (6), was chosen as recommended by Vakirlis et al (Vakirlis et al. 2016) (Supplemental_Fig_S4). The A1 reconstruction was then compared to the other genomes using SynChro (delta values 1-6), and a second ancestral genome (A2) was constructed from A1 and *C. metapsilosis*, with *L. elongisporus* as an outgroup. The best A2 candidate, with the smallest number of chromosomes (15) and conflicts (1) and the highest number of genes (4409) was chosen (Supplemental_Fig_S4). Interchromosomal breaks were identified using pairwise comparison of synteny maps.

### Tagging Cse4

*C. parapsilosis* strains CLIB214 and 90-137 were edited using a tRNA plasmid based CRISPR-Cas9 gene editing system as described by Lombardi et al. (Lombardi et al. 2017, 2019a). Primers gRNA_CSE4_TOP and gRNA_CSE4_BOT were annealed and cloned into pCP-tRNA and 5 µg plasmid was transformed together with 5 µg of a 594 bp synthetic DNA fragment containing a section of the H3 histone variant Cse4 with a 3xHA (hemagglutinin) tag inserted between amino acids 69 and 70, and 250bp homology arms (Integrated DNA Technologies USA, Supplemental_Fig._S1). Transformants were selected on YPD agar supplemented with 200 µg/ml nourseothricin and screened by colony PCR using primers CSE4_N_RT_fw and CSE4_col_inTag_rv. Two clones were sequenced by Sanger sequencing (MWG/Eurofins). Loss of pCP-tRNA was induced by patching transformants onto YPD agar without nourseothricin. For Western blots, protein extracts were prepared from 15 A_600_ units of *C. parapsilosis* 90-137 and two Cse4-HA tagged strains cultured overnight in YPD. Cell pellets were washed in 500 µl water, re-suspended in 500 µl ice-cold extraction buffer (1x PBS, 0.1% Tween 20, 1mM PMFS) and homogenized with glass beads. The protein extract was separated by centrifugation at 10000 rpm at 4. 20 µl protein extracts diluted 1:1 (v/v) with ice-cold 2x Laemmli sample buffer (Sigma-Aldrich) were separated by 12% SDS-PAGE, at 200 V constant voltage for 1 h, and electro-blotted onto nitrocellulose membranes at 100 V for 45 min. Immunoblotting was performed using the mouse epitope tag antibody, Anti-HA.11 (BioLegend), at a 1:1000 dilution in milk/TBS blocking buffer (5 g non-fat dry milk to 100 ml TBS - 100 mM Tris–HCl, pH 7.5, 150 mM NaCl) and HRP-conjugated secondary antibody Anti-mouse IgG (Cell Signaling Technology) at 1:2000 dilution. Immunoblots were detected using the Pierce ECL Western Blotting Substrate (Thermo Scientific) and enhanced chemiluminescence (G:BOX Chemi XRQ, Syngene).

### ChIP-PCR and ChIP-seq

Chromatin immunoprecipitation was carried out as described by Coughlan et al. (Coughlan et al. 2016) from log phase cultures in 200 ml YPD. Control immunoprecipitations were carried out in the absence of the anti-HA antibody (Mock-IP), and from *C. parapsilosis* 90-137 without a tagged Cse4 (CTRL). Dilutions of the protein extracts before immunoprecipitation (Input), and following immunoprecipitation (IP) and mock IP were used to assess binding to CEN1 by PCR amplification, using primers from 5 regions within the predicted CEN1 area, one pair from within the next largest intergenic region on chromosome 1 (chr1:1948277-1955373; to serve as negative control), and a region from within the actin gene *ACT1* (Fig. S1, Table S1). ChIP sequencing was performed by Beijing Genomics Institute (BGI) on the BGISEQ500 platform. Approximately 20 million single-end reads (50 bases) were obtained per sample. ChIP-seq reads were mapped to the genome of *C. parapsilosis* CDC317 (Butler et al. 2009) using the aln/samse algorithm from BWA v0.7.17-r1188 (Li and Durbin 2010), with default parameters. Mapped reads were sorted and indexed with Samtools v 1.9 (Li et al. 2009) and the read coverage across the genome was computed using Bedtools v2.27.1 (Quinlan and Hall 2010). Genome coverage files were changed into bigwig format using bedGraphToBigWig v4 (Kent et al. 2010) and loaded into Integrated Genomics Viewer (Thorvaldsdóttir et al. 2013) for visualization.

### minION sequencing

One derivative of *C. parapsilosis* 90-137 containing Cse4-HA was sequenced using the minION device from Oxford Nanopore Technologies (ONT). DNA was extracted using the MagJET Genomic DNA Kit #K2721 from ThermoScientific. Libraries were prepared with the Rapid Sequencing Kit (RSK-SQK004) from ONT and sequenced on a minION flow cell (FLO-MIN106), yielding 30X coverage. Basecalling was performed using Guppy v2.3.7+e041753. Read length and quality was assessed using NanoPlot v1.23.1 (De Coster et al. 2018). NanoFilt v2.3.0 (De Coster et al. 2018) was used to remove reads with a quality score of less than 7, and the assembly was constructed using canu v1.8 (Koren et al. 2017) with options genomeSize=13030174 (to specify the genome size) and -nanopore-raw (for ONT data). Nanopolish v0.11.1 (Loman et al. 2015) was used to improve the consensus accuracy of the assembly and the sequence quality was further improved by incorporating the BGISEQ data from the “input” sample of the ChIP-seq experiment using Pilon v1.23 (Walker et al. 2014). The assembly quality was assessed with Quast v4.6.1 (Gurevich et al. 2013). Circoletto and Circos v0.69 (Darzentas 2010; Krzywinski et al. 2009) were used to visualise alignments between the *C. parapsilosis* CDC317 reference genome and *C. parapsilosis* 90-137/Cse4-HA assembly. Contigs of less than 1kb and/or contigs mapping to the mitochondrial genome were removed from the assembly.

### Data access

All data generated for this project has been deposited in GenBank under the NCBI BioProject ID PRJNA563885.

## ACKNOWLEDGEMENTS

This work was supported by Science Foundation Ireland (12/IA/1343 (GB) and 13/IA/1910 (KW), https://www.sfi.ie), The Wellcome Trust (102406/Z/13/Z and 109167/Z/15/Z, https://wellcome.ac.uk) and the European Research Council (789341) (KW).

## Supplemental material

**Fig. S1.**
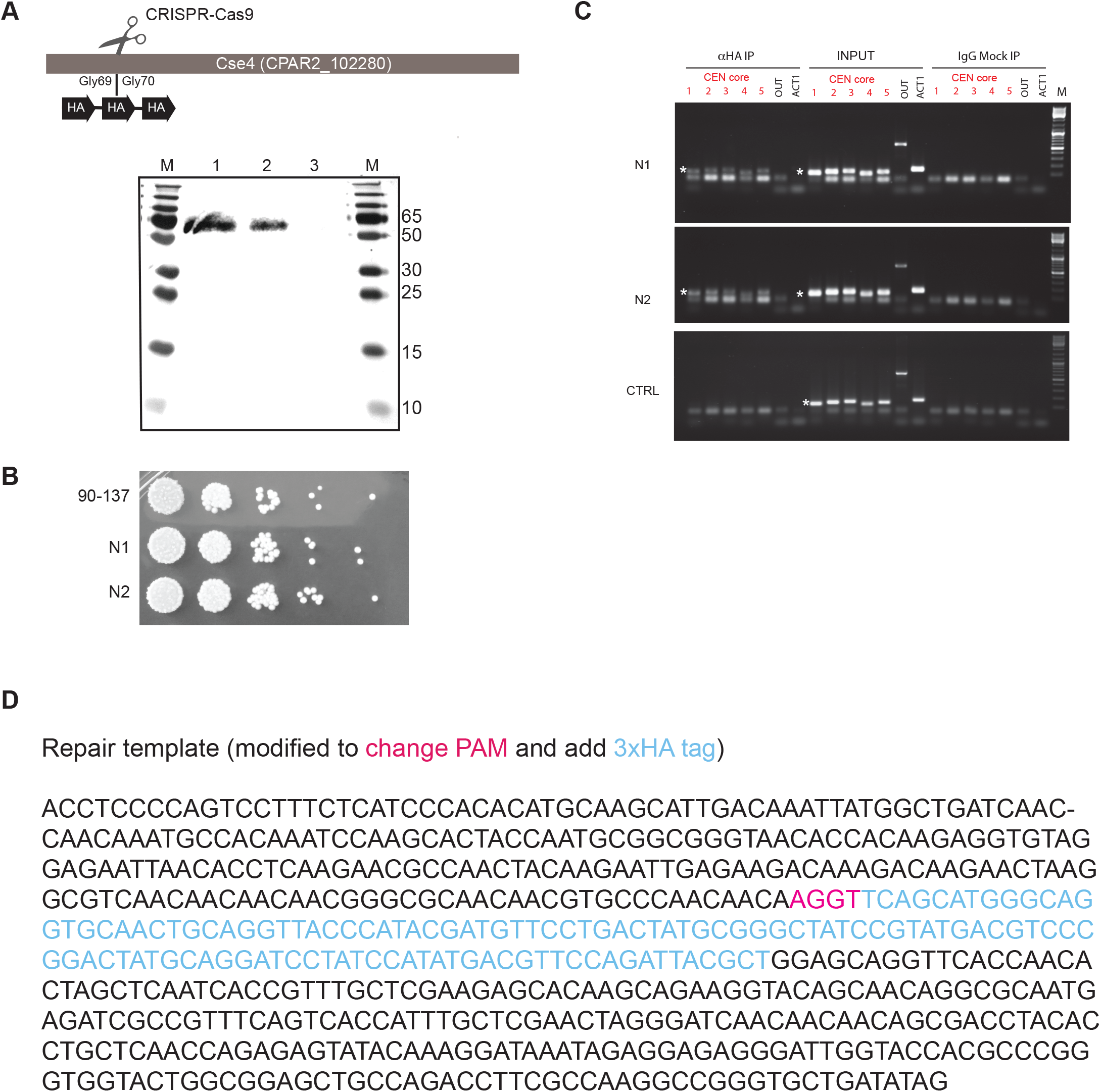
Expression of Cse4-HA and ChIP-PCR. A. Expression of Cse4-HA in *C. parapsilosis* 90-137 was detected by Western blot using anti-HA antibody. Lane M: PageRuler Plus Prestained Protein ladder (Thermo Scientifics). Sizes are shown in kD. Lanes 1 and 2: protein extract from two independent Cse4-HA tagged derivatives of *C. parapsilosis* 90-137 Lane 3: protein extract from untagged *C. parapsilosis* 90-137. The protein size is expected to be 49.4 kD. B. Tagging Cse4 does not interfere with growth. The tag was introduced at a position that was predicted to be unlikely to interfere with the function of Cse4. Two Cse4-tagged derivatives of *C. parapsilosis* 90-137 (N1 and N2) grow as well as the parental strain *C. C. parapsilosis* 90-137 on YPD. Serial dilutions of cells were spotted on YPD agar and grown for 48h at 30° C. ChIP-PCR from *C. parapsilosis* CEN1. N1 and N2 are Cse4-tagged derivatives, and CTRL is the untagged strain. PCR amplification was carried out using 5 pairs of primers from within the core region of CEN1 shown in red (Table S1), one pair from an adjacent intergenic region on Chromosome 1 (OUT) and one pair from *ACT1*. IP = HA immunoprecipitation with anti-HA; INPUT = protein extract before immunoprecipitation; mock IP = no anti-HA antibody used. The target PCR products in the core region are marked with a white asterisk. Some smaller non-specific products were also obtained. All primer pairs from within CEN1 amplified the expected size fragments from total chromatin (Input) from untagged *C. parapsilosis* 90-137 and from two Cse4-tagged strains. Some non-specific PCR products were also amplified. The CEN1-specific primers also amplified sequences from anti-HA chromatin immunoprecipitates in the tagged strains (N1 and N2), but not in the control untagged strain (CTRL). The OUT and *ACT1* primers amplified products from the input samples only. Cse4 therefore localizes to the proposed CEN1. D. Sequence of repair template used to introduce HA tags.

**Fig. S2.**
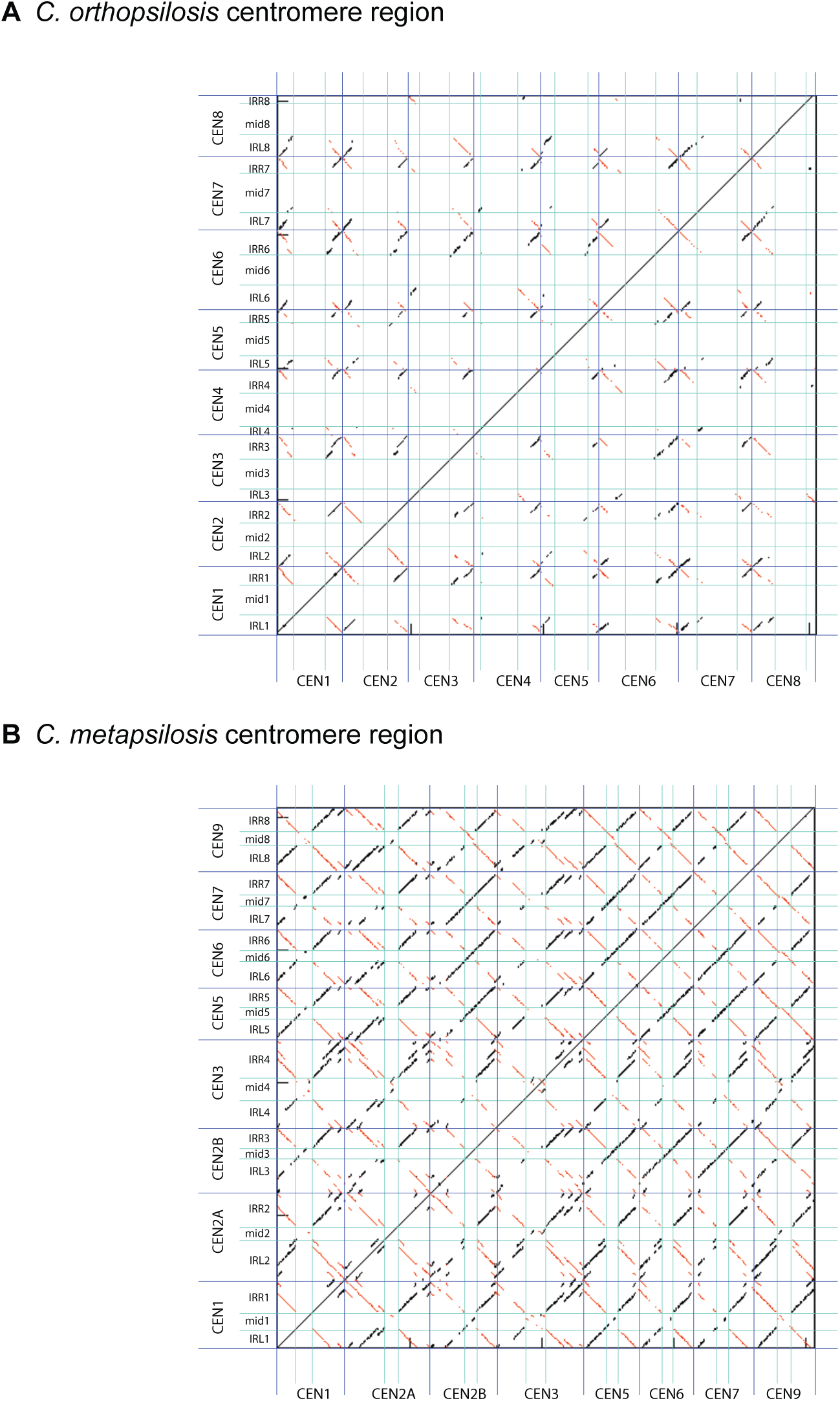
Dot matrix plots showing sequence conservation around *C. orthopsilosis* (A) and *C. metapsilosis* centromeres (B). Centromeres are delineated by dark blue lines. Inverted repeats (Right, IRR and left, IRL) are separated with cyan lines. Each dot represents identity of 25-bp. Inverted sequences are shown in red, and direct repeats in black.

**Fig. S3.**
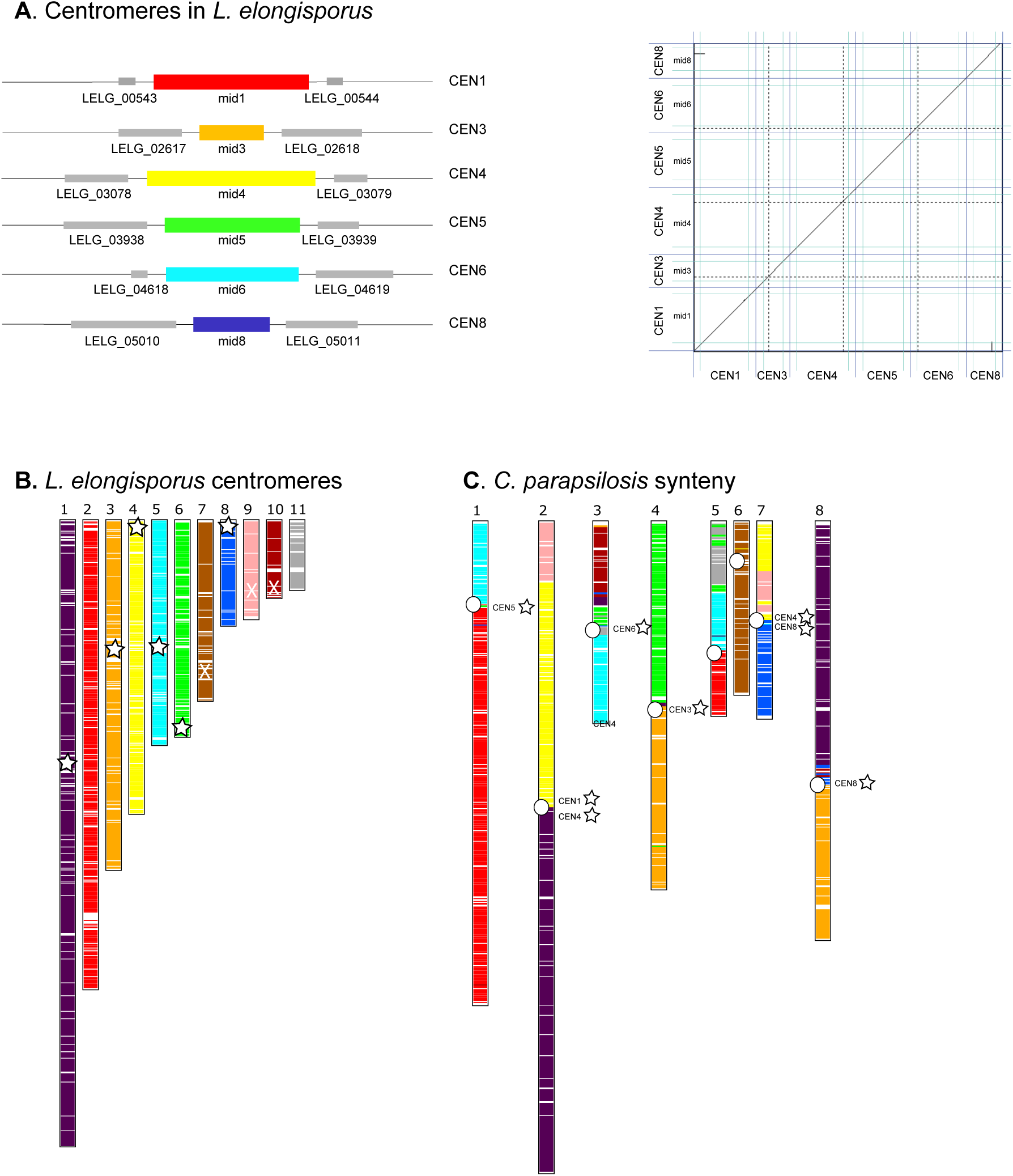
Rearrangements at putative centromeres in *L. elongisporus*. Synteny relationship between *C. parapsilosis* and *L. elongisporus* identified using SynChro. A. Location of hits on *L. elongisporus* chromosomes. The approximate location of the putative *L. elongisporus* centromeres are indicated with white stars. Three proposed candidates (marked with white X’s) are unlikely to represent centromeres because they are either adjacent to the rDNA (scaffold 9), or on regions that are strongly transcribed (scaffolds 7 and 10). The other proposed centromeres are unique, when only the untranscribed regions are included (Donovan et al. 2016). B. *C. parapsilosis* chromosomes, colored with respect to the RBHs from *L. elongisporus*. The location of the putative *C. parapsilosis* centromeres are indicated with a white circle. The location of syntenic *L. elongisporus* centromeres (where identified) are indicated by name and with a white star.

**Fig S4.**
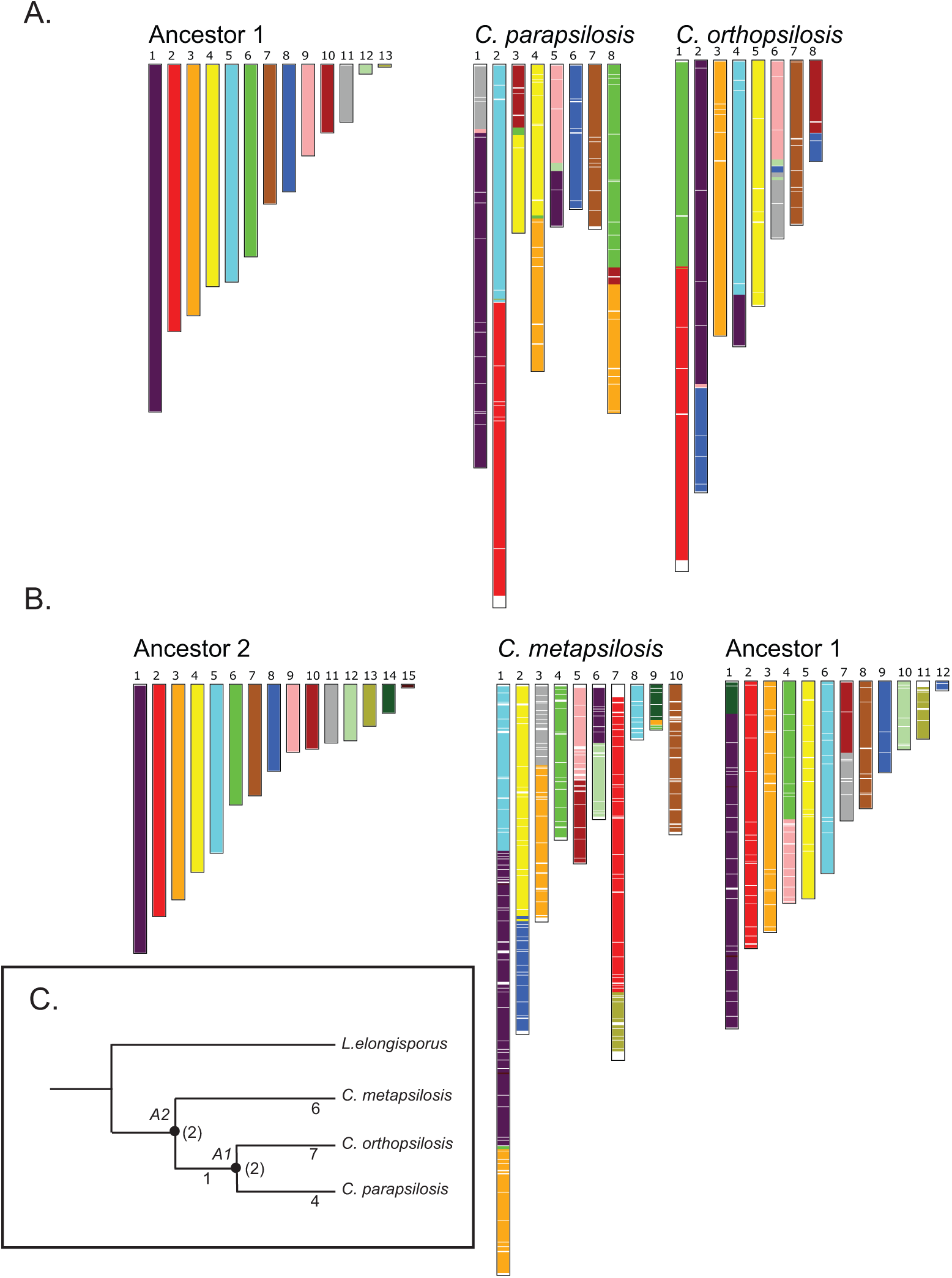
Ancestral reconstruction. A. The genome structure of the ancestor of *C. parapsilosis* and *C. orthopsilosis* (Ancestor 1, A1) was inferred using AnChro (Vakirlis et al. 2016). The syntenic relationship between A1 and C. parapsilosis or C. orthopsilosis (determined using SynChro) is shown (see Fig. 4). B. The genome structure of the ancestor of A1 and *C. metapsilosis* (Ancestor 2, A2) was inferred as in (A). C. Interchromosomal breaks were identified by pairwise comparisons of synteny maps from SynChro. Thirteen breaks were identified between *C. parapsilosis* and *C. orthopsilosis*. By comparing with A1, 7 were placed on the C. orthopsilosis branch, and 4 on the C. parapsilosis branch. Two (shown in parentheses) could not be placed. Similar comparisons are shown with Ancestor A2. The phylogenetic relationship between the species is taken from Pryszcz et al (2015).

**Table S1.**
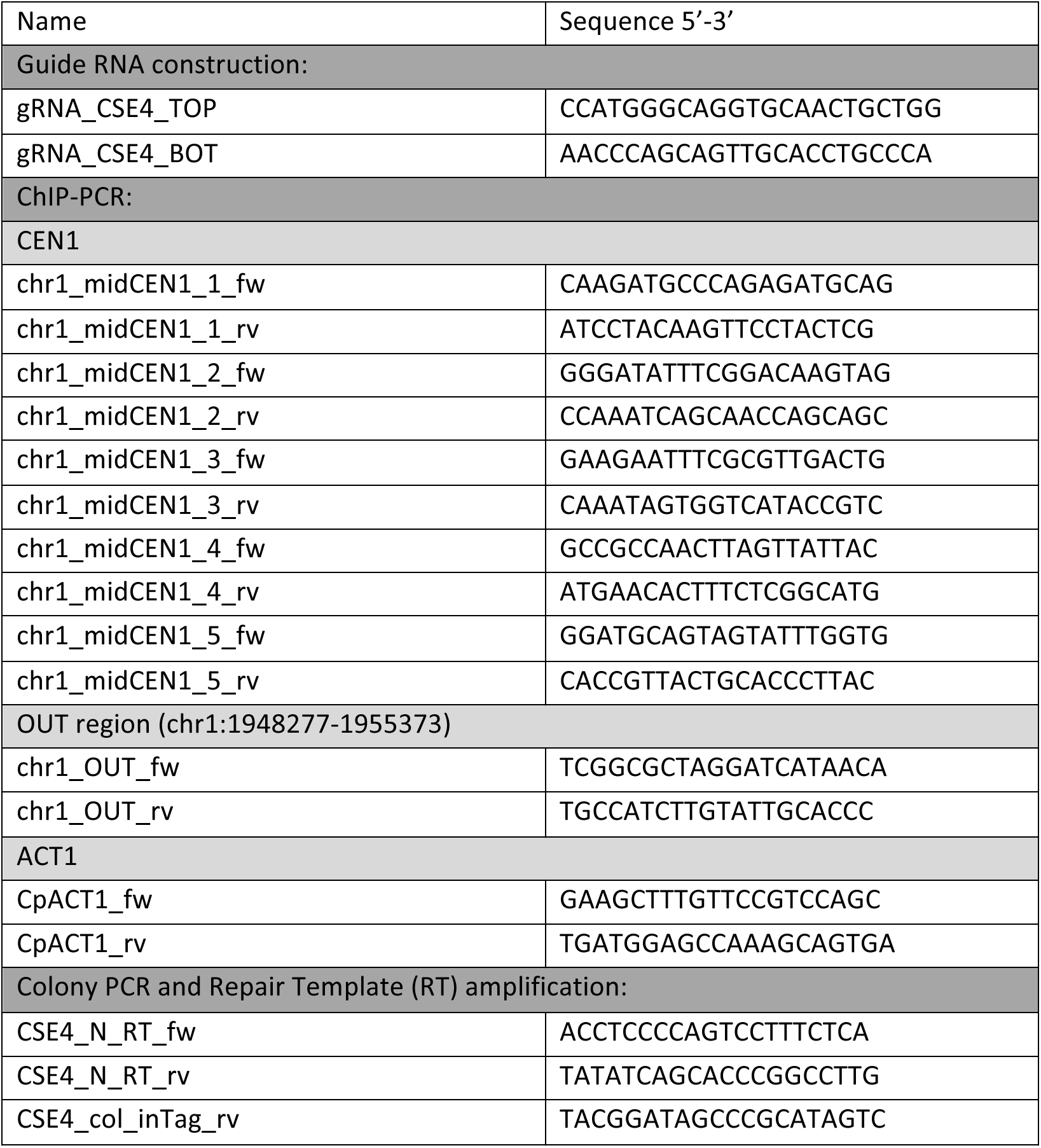
List of primers used.

**Table S2.**
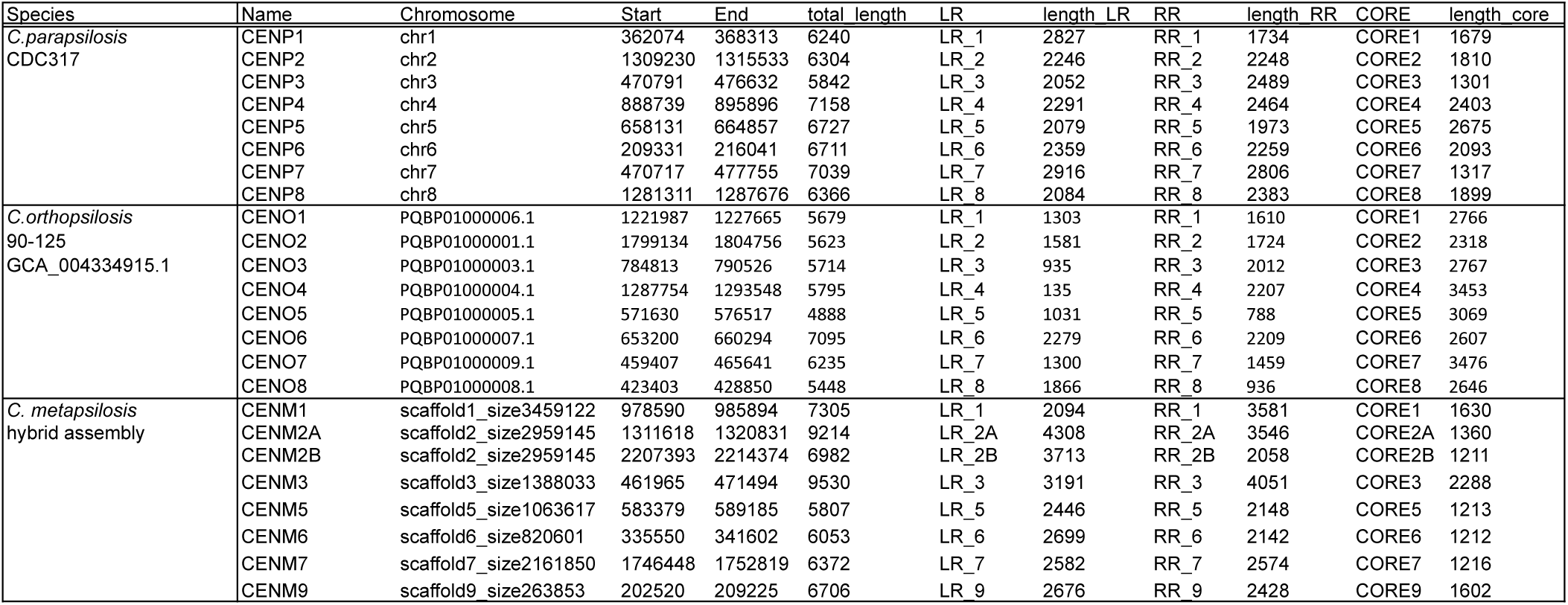
Coordinates of centromeres in *C. parapsilosis, C. orthopsilosis* and *C. metapsilosis*. Numbers for *C. parapsilosis* (CDC317) and *C. metapsilosis* (hybrid) refer to the current assembly for each, available at CGOB (Fitzpatrick et al. 2010). The *C. orthopsilosis* locations refer to the minION assembly of *C. orthopsilosis* 90-125 from Lombardi et al (Lombardi et al. 2019b).

| Species | Name | Chromosome | Start | End | total length | LR | length LR | RR | length RR | CORE | length core |
| --- | --- | --- | --- | --- | --- | --- | --- | --- | --- | --- | --- |
| *C.parapsilosis*<br>CDC317 | CENP1 | chr1 | 362074 | 368313 | 6240 | LR_1 | 2827 | RR_1 | 1734 | CORE1 | 1679 |
|  | CENP2 | chr2 | 1309230 | 1315533 | 6304 | LR_2 | 2246 | RR_2 | 2248 | CORE2 | 1810 |
|  | CENP3 | chr3 | 470791 | 476632 | 5842 | LR_3 | 2052 | RR_3 | 2489 | CORE3 | 1301 |
|  | CENP4 | chr4 | 888739 | 895896 | 7158 | LR_4 | 2291 | RR_4 | 2464 | CORE4 | 2403 |
|  | CENP5 | chr5 | 658131 | 664857 | 6727 | LR_5 | 2079 | RR_5 | 1973 | CORE5 | 2675 |
|  | CENP6 | chr6 | 209331 | 216041 | 6711 | LR_6 | 2359 | RR_6 | 2259 | CORE6 | 2093 |
|  | CENP7 | chr7 | 470717 | 477755 | 7039 | LR_7 | 2916 | RR_7 | 2806 | CORE7 | 1317 |
|  | CENP8 | chr8 | 1281311 | 1287676 | 6366 | LR_8 | 2084 | RR_8 | 2383 | CORE8 | 1899 |
| *C.orthopsilosis*<br>90-125<br>GCA_004334915.1 | CENO1 | PQBP01000006.1 | 1221987 | 1227665 | 5679 | LR_1 | 1303 | RR_1 | 1610 | CORE1 | 2766 |
|  | CENO2 | PQBP01000001.1 | 1799134 | 1804756 | 5623 | LR_2 | 1581 | RR_2 | 1724 | CORE2 | 2318 |
|  | CENO3 | PQBP01000003.1 | 784813 | 790526 | 5714 | LR_3 | 935 | RR_3 | 2012 | CORE3 | 2767 |
|  | CENO4 | PQBP01000004.1 | 1287754 | 1293548 | 5795 | LR_4 | 135 | RR_4 | 2207 | CORE4 | 3453 |
|  | CENO5 | PQBP01000005.1 | 571630 | 576517 | 4888 | LR_5 | 1031 | RR_5 | 788 | CORE5 | 3069 |
|  | CENO6 | PQBP01000007.1 | 653200 | 660294 | 7095 | LR_6 | 2279 | RR_6 | 2209 | CORE6 | 2607 |
|  | CENO7 | PQBP01000009.1 | 459407 | 465641 | 6235 | LR_7 | 1300 | RR_7 | 1459 | CORE7 | 3476 |
|  | CENO8 | PQBP01000008.1 | 423403 | 428850 | 5448 | LR_8 | 1866 | RR_8 | 936 | CORE8 | 2646 |
| *C. metapsilosis*<br>hybrid assembly | CENM1 | scaffold1_size3459122 | 978590 | 985894 | 7305 | LR_1 | 2094 | RR_1 | 3581 | CORE1 | 1630 |
|  | CENM2A | scaffold2_size2959145 | 1311618 | 1320831 | 9214 | LR_2A | 4308 | RR_2A | 3546 | CORE2A | 1360 |
|  | CENM2B | scaffold2_size2959145 | 2207393 | 2214374 | 6982 | LR_2B | 3713 | RR_2B | 2058 | CORE2B | 1211 |
|  | CENM3 | scaffold3_size1388033 | 461965 | 471494 | 9530 | LR_3 | 3191 | RR_3 | 4051 | CORE3 | 2288 |
|  | CENM5 | scaffold5_size1063617 | 583379 | 589185 | 5807 | LR_5 | 2446 | RR_5 | 2148 | CORE5 | 1213 |
|  | CENM6 | scaffold6_size820601 | 335550 | 341602 | 6053 | LR_6 | 2699 | RR_6 | 2142 | CORE6 | 1212 |
|  | CENM7 | scaffold7_size2161850 | 1746448 | 1752819 | 6372 | LR_7 | 2582 | RR_7 | 2574 | CORE7 | 1216 |
|  | CENM9 | scaffold9_size263853 | 202520 | 209225 | 6706 | LR_9 | 2676 | RR_9 | 2428 | CORE9 | 1602 |

